# Rapid characterization of complex genomic regions using Cas9 enrichment and Nanopore sequencing

**DOI:** 10.1101/2021.03.11.434935

**Authors:** Jesse Bruijnesteijn, Marit van der Wiel, Natasja G. de Groot, Ronald E. Bontrop

## Abstract

Long-read sequencing approaches have considerably improved the quality and contiguity of genome assemblies. Such platforms bear the potential to resolve even extremely complex regions, such as multigenic families and repetitive stretches of DNA. Deep sequencing coverage, however, is required to overcome low nucleotide accuracy, especially in regions with high homopolymer density, copy number variation, and sequence similarity, such as the *MHC* and *KIR* gene clusters of the immune system. Therefore, we have adapted a targeted enrichment protocol in combination with long-read sequencing to efficiently annotate complex genomic regions. Using Cas9 endonuclease activity, segments of the complex *KIR* gene cluster were enriched and sequenced on an Oxford Nanopore Technologies platform. This provided sufficient coverage to accurately resolve and phase highly complex *KIR* haplotypes. Our strategy facilitates rapid characterization of large and complex multigenic regions, including its epigenetic footprint, in multiple species, even in the absence of a reference genome.

## Introduction

Repetitive regions are difficult to resolve using short-read sequencing approaches, and often remain registered for years as incomplete gaps in draft genomes (Chinwalla et al. 2002; International Human Genome Sequencing 2004; Lindblad-Toh et al. 2005; Waterson et al. 2005; Gibbs et al. 2007; Marques-Bonet et al. 2009; Alkan et al. 2011). These complex stretches often involve transposable elements, microsatellites, and multi-copy gene clusters, the latter of which is represented by multiple gene families that encode essential components of the immune system (Thomma et al. 2016; Peona et al. 2019). For example, the major histocompatibility complex (MHC) genes, known in humans as human leukocyte antigens (HLA) genes, are considered the most polymorphic gene cluster. The *MHC* genes co-evolved with their ligands of the killer cell immunoglobulin-like receptor (KIR) gene family, which also features striking levels of complexity.

The fundamental limitations to characterizing these complex regions by short-read sequencing strategies are potentially overcome by third-generation techniques that generate high yields of long reads (Eid et al. 2009; Jain et al. 2015). Oxford Nanopore sequencing may produce reads far above 100 kb by recording changes in the electrical current as nucleotides pass through synthetic nanopores. The data quality and throughput of nanopore sequencing is improving rapidly, and has allowed the *de novo* assembly of multiple human genomes (Jain et al. 2018; Cai et al. 2020; Shafin et al. 2020). These genome assemblies contiguously span multigenic clusters, such as the *MHC* and *KIR* gene regions, but correct annotation is hampered by the relatively low coverage, which precludes at this stage an accurate allele level resolution. Considering the important role of different multi-copy gene families in health and disease, a cost-efficient and high-resolution characterization approach regarding these types of regions is urgent.

Instead of whole genome sequencing, specific genes and regions might be enriched during library preparation. For instance, the *MHC class II DRB* gene region was enriched by long-ranged PCR, and characterized using a hybrid sequencing approach that combined Illumina and Oxford Nanopore platforms (Fuselli et al. 2018). Amplification steps, however, might introduce nucleotide errors during synthesis and, in addition, erase all epigenetic footprints. An amplification-free enrichment technique involves Cas9-mediated targeting of chromosome segments and nanopore sequencing (Gabrieli et al. 2018; Giesselmann et al. 2019; Gilpatrick et al. 2020; Stangl et al. 2020; Watson et al. 2020). The Cas9 endonuclease activity may specifically excise genomic regions of interest that are subsequently ligated to nanopore adapters. This allows the direct sequencing of genomic segments while avoiding error prone DNA synthesis and maintaining epigenetic modifications. Efficient and specific enrichment using this approach has been demonstrated for single genes, including several cancer-related fusion genes (Stangl et al. 2020), but an application for multigenic regions is absent in the literature.

In this study, we adapted the Cas9-mediated enrichment method to resolve complex multigenic regions, and validated this approach by the targeted characterization of complex *KIR* gene clusters in two different primate species. We focused on the *KIR* region in humans, which has been thoroughly characterized at the genomic level, and is important, for instance, in AIDS susceptibility and transplantation biology (Farag et al. 2006; Bashirova et al. 2011). Rhesus macaques, however, represent an important species in preclinical health research concerning, for example, COVID-19 and AIDS (Hatziioannou et al. 2009; Estes et al. 2018; Yu et al. 2020), but the physical location of the *KIR* genes is poorly understood. The *KIR* receptor family is involved in the regulation of NK cell activity, and comprises activating and inhibitory members that may recognize particular epitopes on MHC class I molecules. A comprehensive nomenclature system distinguishes the variety of KIR receptors, and reflects the number of extracellular domains (KIR1D, KIR2D, KIR3D) and the length of the cytoplasmic tail (long, L; short, S) (Marsh et al. 2003; Bruijnesteijn et al. 2020b). Subsequent numbering defines structurally similar but phylogenetically distinct genes (e.g., *KIR2DL1*), whereas three additional digits distinguish allotypes (e.g., *KIR2DL1*001*). In humans, a total of 17 *KIR* genes are defined, 1110 alleles of which are documented (IPD-KIR, release 2.9.0) (Maccari et al. 2020).

The *KIR* gene repertoire is shaped by abundant tandem duplications, deletions, and chromosomal recombination events, and exceeds the plasticity of the *MHC* gene cluster (Guethlein et al. 2007; Pyo et al. 2013; de Groot et al. 2015). The *KIR* genes are 10 to 15 kb long, and are arranged in a head-to-tail manner, separated by intergenic regions of approximately 2 kb. Sequence similarity characterizes the genetic cluster, with any two *KIR* genes sharing 80%–90% homology, and allelic variants of a certain gene tend to be over 98% similar. The KIR haplotypes, defined as a segregating unit of genes located on a single chromosome, distinguish a centromeric and telomeric segment, which display diverse configurations and extensive copy number variation. For this study, we resolved six human *KIR* haplotypes, derived from three randomly selected human donors, at an allele-level resolution.

The continuous evolution of the *KIR* gene system is reflected by the genomic diversification at a species, population, and individual level. To validate our concept, we enriched and assembled KIR haplotypes in rhesus macaques. So far, only two completely sequenced rhesus macaque *KIR* haplotypes have been documented, but previous transcriptome and segregation studies indicate extensive variation (Sambrook et al. 2005; Bimber et al. 2008; Bruijnesteijn et al. 2018; Dutcher 2018; Bruijnesteijn et al. 2020a). Annotation of this complex immune region is a difficult enterprise, which is reflected by a poorly annotated *KIR* region in the rhesus macaque reference genome (Mmul_10) (Warren et al. 2020). Our genomic characterization of rhesus macaque *KIR* haplotypes demonstrated the rapid construction of complex multigenic haplotypes, even in the absence of reference sequences. Hence, adaption of this technique would allow a speedy and cost-efficient characterization of other multigenic regions, from which whole genome assemblies and clinical implications might benefit.

## Results

### A ‘tiling’ approach to enrich complex multigenic clusters without amplification

The characterization of large and repetitive immune regions requires the generation and sequencing of genomic DNA (gDNA) fragments that share overlaps. Allelic variation in these overlaps allows the phasing of haplotypes. To achieve this goal, dephosphorylated high molecular weight (HMW) gDNA needs to be cleaved, using sets of CRISPR RNAs (crRNA) in complex with Cas9 endonuclease (Fig. S1). These crRNAs are designed to target conserved stretches that are shared by members of a multigenic family. This approach will allow generic enrichment. Only at the terminus of the cleaved target sites is a phosphate group available, which is utilized for dA-tailing and subsequent ligation to Nanopore sequencing adaptors. This ‘tiling’ approach facilitates the selective enrichment of large overlapping DNA segments, and allows the subsequent sequencing of polymorphic multigenic regions without the need for amplification.

### Enrichment of complex KIR regions

To validate our approach, the *KIR* gene regions in humans and rhesus macaques were enriched and characterized. The nature of *KIR* gene complexity required the design of species-specific sets of crRNAs to enrich complete genomic clusters (Sup. Table 1 & 2). In humans, the presence of four framework genes with distinctive physical locations marks the centromeric (*KIR3DL3* and *KIR3DP1*) and telomeric (*KIR2DL4* and *KIR3DL2*) haplotype segments (Fig. 1). Expansion and contraction of both regions resulted in haplotypes that contain 9 to 14 *KIR* genes, including two pseudogenes (*KIR2DP1* and *KIR3DP1*). Important to note, however, is that some genes that are present on a given *KIR* haplotype may be absent from another. Human *KIR* haplotypes can be roughly categorized based on their gene content, into those with more an inhibitory (group A) or an activating (group B) gene profile (Uhrberg et al. 1997). Recombination events, possibly owing to the high transposon density, might rearrange haplotype organizations (Fig. 1). To determine the high content variability of this genetic cluster, 35 generic crRNAs were designed to target the differential presence of human *KIR* genes that may be encountered on a haplotype, whereas 12 crRNA were specific for one particular framework gene (Sup. Table 1). In addition, seven crRNA were included to target the genes that flank the *KIR* gene cluster (*LILR* and *FcAR*), in order to define both ends of the *KIR* haplotype.

**Figure 1.**
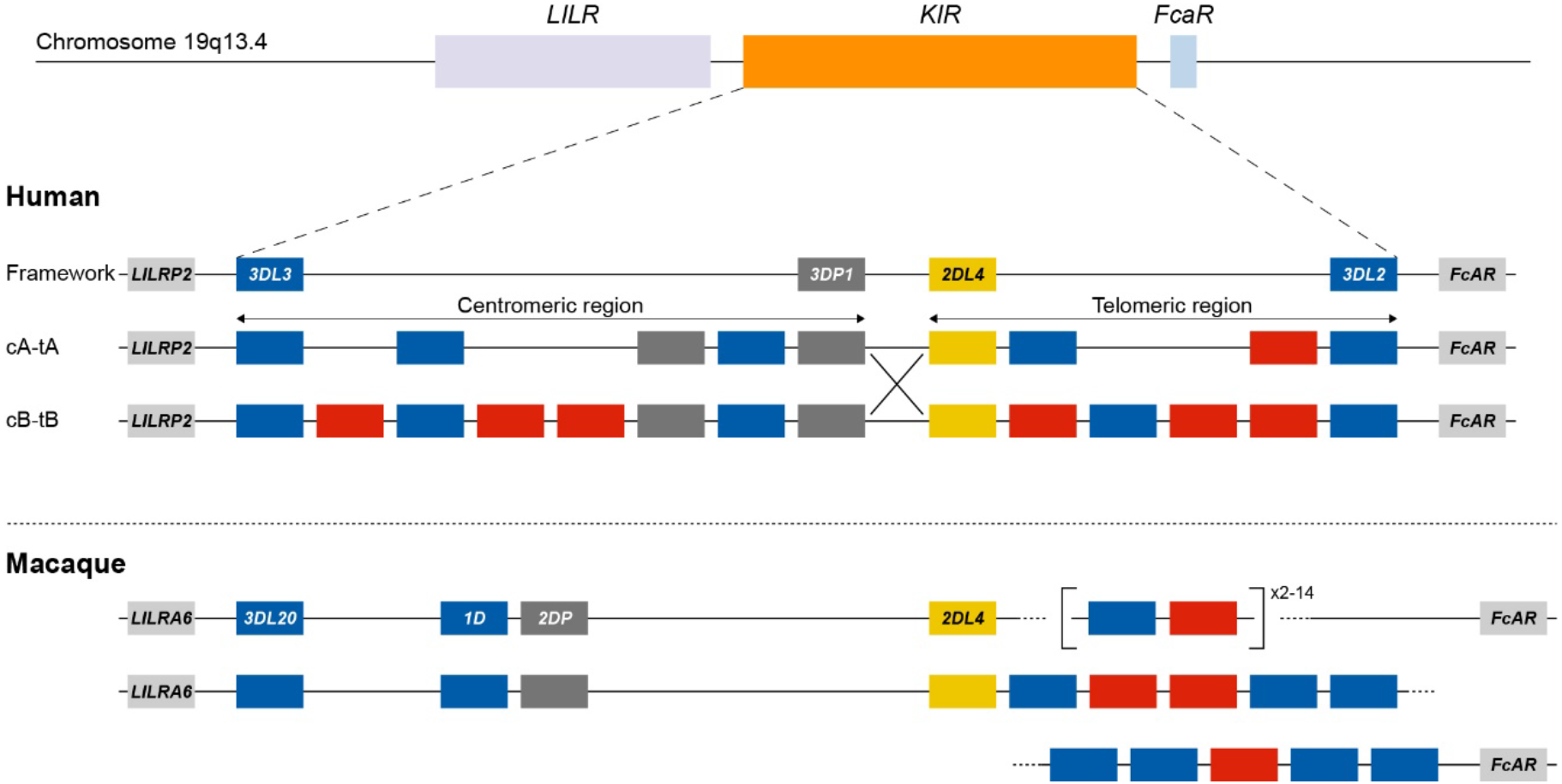
*KIR* haplotype configurations in humans and rhesus macaques. The *KIR* gene family maps between the *LILR* cluster and the *FcAR* gene. In humans, four genes are considered to reflect a framework reference haplotype, with *KIR3DL3* and *KIR3DP1* defining the centromeric region (c), and *KIR2DL4* and *KIR3DL2* bounding the telomeric region (t). In the human population, both the c and t regions may feature different gene contents. The red and blue boxes depict activating or inhibitory receptors, respectively. Pseudogenes have been indicated as well (grey). The only homologous gene shared in humans and macaques is *KIR2DL4* (yellow). Human *KIR* haplotypes are categorized based on a more inhibitory (group A) or activating (group B) gene profile, and reflect different segment configurations (e.g., cA-tA and cB-tB) [31]. Chromosomal recombination events might shuffle haplotype segments (e.g., cA-tB defined by the large X). In rhesus macaques, the framework is represented by *KIR3DL20* at the centromeric region. The telomeric region is characterized by highly variable gene expansions and contractions. In contrast to the situation for humans, physical maps of rhesus macaque *KIR* haplotypes are virtually absent from the literature. For that reason, the lower panel represents a hypothetical configuration as the proposed situation at the start of this study.

The *KIR* haplotypes in rhesus macaques display even more content diversity, with 4 to 17 KIR transcripts encoded as defined by segregation studies (Bruijnesteijn et al. 2018; Bruijnesteijn et al. 2020a). The only framework gene present on all haplotypes is *KIR3DL20*, which marks the centromeric region (Fig. 1). Only two other *KIR* genes are differentially located within this haplotype segment (*KIR1D* and *KIR2DP*). The only macaque KIR ortholog that is shared with humans, *KIR2DL04*, is present on approximately 80% of the telomeric haplotype segments. This haplotype region is further characterized by a differential number of *KIR* genes in highly diverse frequencies. To enrich the macaque *KIR* cluster, we designed 24 generic crRNAs that target the variable gene tandem, and complemented those with crRNAs specific for *KIR1D*, *KIR2DL04*, and *KIR3DL20*, and the flanking genes, *LILRA6* and *FcAR* (Sup. Table 2).

### Assembly of human KIR haplotypes

For this communication, the human target region was defined at the start of exon 5 of *LILRP2* to the end of exon 1 of *FcAR*, thereby comprising the complete *KIR* haplotype (Fig. 1). On the human reference genome (HG38), this target region spans approximately 161 kb and contains nine *KIR* genes, one of which encodes an activating KIR (group A haplotype). To account for the variability of *KIR* haplotype configurations, four additional reference sequences, which were assembled by Fosmid sequencing and reflected different *KIR* genotypes, were included in our panel (Roe et al. 2017). In total, five reference *KIR* regions were used to assemble captured reads and to determine sequence accuracy.

For each of the three randomly selected human individuals, two Nanopore flowcells were loaded with MHW gDNA samples that were enriched either for fragments that comprise a complete *KIR* gene or fragments that connect and distinguish neighboring *KIR* genes. An average of 1.26 million Nanopore reads were yielded per flowcell (Table 1). Of these reads, 4.2% to 19.3% had a length of 7,000 kb or longer, which is required to connect and distinguish neighboring *KIR* genes. The percentage of size-selected reads that mapped to the target region ranged from 1.6% to 2.5% (± 4,015 reads), which provided a median coverage ranging from 269 to 323X. The enrichment factor, which reflects the efficiency of the targeted enrichment, ranged from 215 to 394X. The length of the consensus sequences that were generated from the on-target reads ranged from 6.5 to 29.7 kb, and their assembly covered complete reference *KIR* haplotype configurations (Fig. 2A, 2B, and 2C). The consensus accuracy compared to the reference sequences ranged from 96.7% to 99.9%.

**Table 1:** Overview of the total reads, read length, on-target hits, and the coverage of the target region in human samples.

|  | Gene fragments (flowcell 1) |  |  |
| --- | --- | --- | --- |
|  | #1 | #2 | #3 |
| Total # reads | 789705 | 1175600 | 1413919 |
| Total # reads > 7.000 bp | 57830 | 49535 | 59575 |
| Percentage reads > 7.000 bp | 7.3 | 4.2 | 4.2 |
| Average read length (bp) | 2603 | 2114 | 2092 |
| Average read length (bp) of reads > 7.000 bp | 13410 | 10895 | 13317 |
|  | Gene to gene fragments (flowcell 2) |  |  |
|  | #1 | #2 | #3 |
| Total # reads | 1060000 | 1078502 | 2012903 |
| Total # reads > 7.000 bp | 204797 | 152372 | 108688 |
| Percentage reads > 7.000 bp (%) | 19.3 | 14.1 | 5.4 |
| Average read length (bp) | 5062 | 3728 | 2021 |
| Average read length (bp) of reads > 7.000 bp | 17206 | 13327 | 13455 |
|  | Total reads (flowcells 1 and 2) |  |  |
|  | #1 | #2 | #3 |
| On-target reads (> 7.000 bp reads) | 4470 | 3308 | 4268 |
| Percentage on-target reads | 1.7 | 1.6 | 2.5 |
| Mean coverage target region | 323 | 269 | 315 |
| Enrichment factor | 215 | 299 | 394 |

**Figure 2.**
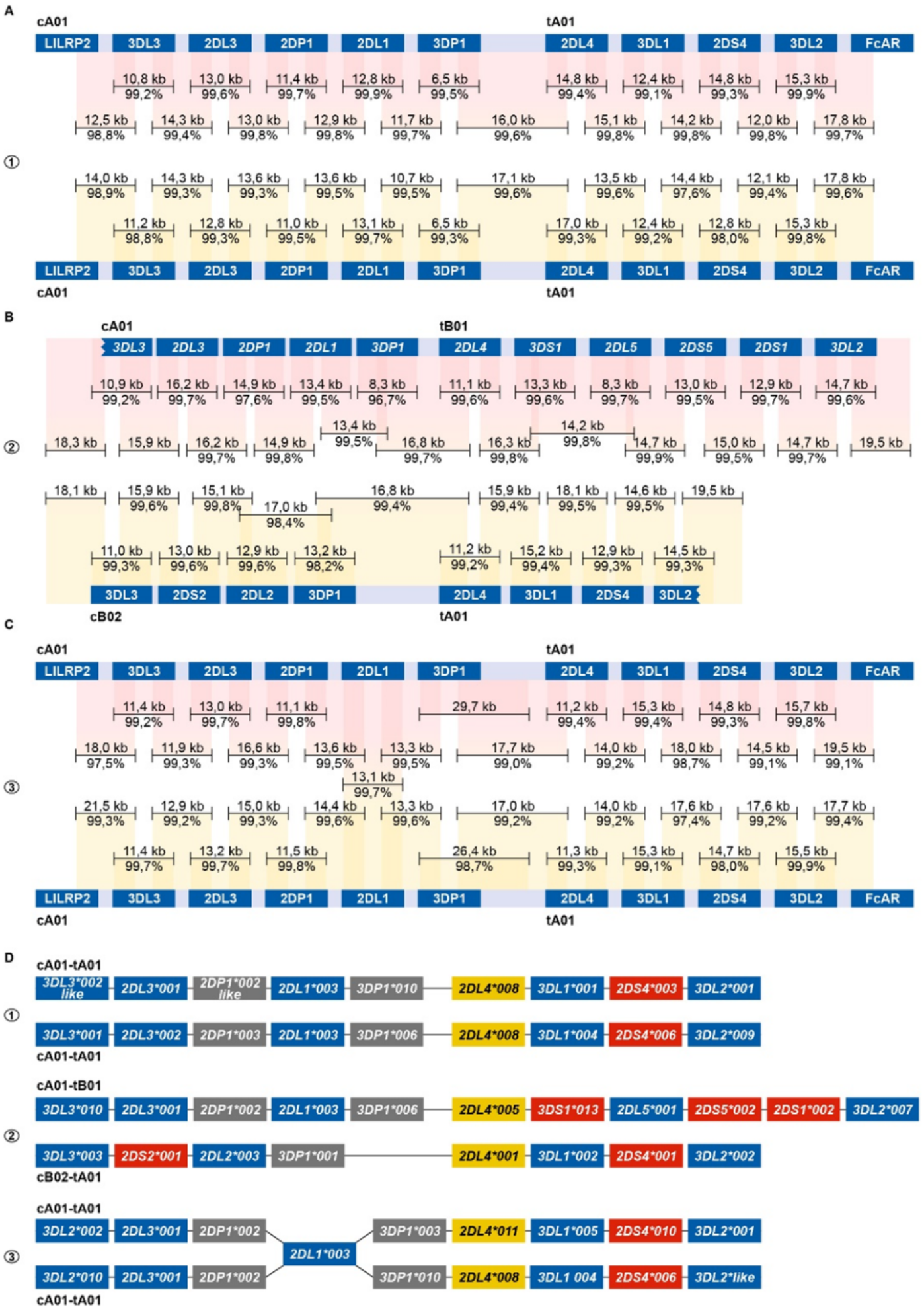
Enrichment and phasing of human *KIR* haplotypes. Human donors 1, 2, and 3 were randomly selected, and displayed one heterozygous (#2) and two homozygous (#1, #3) *KIR* haplotype configurations. Consensus sequences (black bars) that covered *KIR* genes from start to end, or that comprised segments from neighbouring *KIR* genes, were mapped to corresponding haplotype references derived from the human reference genome (HG38) or from previous Fosmid sequencing studies (Roe et al. 2017). The consensus sequence length and accuracy are indicated. Colored columns indicate the mapped regions, in which a darker color illustrates overlapping consensus sequences. *KIR* haplotypes were completely phased in two individuals (A, B). The third individual shared an identical *KIR2DL1* gene on both haplotypes, which located an extended SNP desert (25 kb; Fig. S2) (C). *KIR* haplotypes of this individual were phased at all genes at an allele level resolution, except for the *KIR2DL1* gene. Four differential KIR region configurations from three randomly selected donors were resolved. Based on allelic polymorphism, each of these haplotypes appears to be unique (D). In addition, three new variants of documented alleles were defined, which are indicated with “like”.

The centromeric and telomeric regions were completely assembled and grouped into four different segment configurations (cA01, cB02, tA01, tB01). Phasing of complete KIR haplotypes was achieved in two individuals, including a homozygous haplotype configuration using allele level resolution (Fig. 2A, 2B and 2D). This result demonstrates the resolution power of our approach. For another homozygous setting, *KIR* haplotypes were phased for all genes, except for *KIR2DL1* (Fig. 2C). This gene was identical on both haplotypes, and defined a SNP desert (25 kb) that hampered complete phasing (Fig. S2). The largest completely assembled and phased haplotype comprised 11 *KIR* genes and covered a total of approximately 176 kb (Fig. 2B).

### Assembly of rhesus macaque KIR haplotypes without reference genome

The successful deciphering of highly variable rhesus macaque *KIR* haplotypes validated our enrichment and characterization approach further, even in the absence of genomic reference sequences. The haplotype content was initially determined by previous transcriptome and segregation studies (Bruijnesteijn et al. 2018; Bruijnesteijn et al. 2020a), but the physical location of *KIR* genes remained elusive. The target region was defined at the start of exon 7 in *LILRA6* to the end of exon 2 in *FcAR*, thereby comprising the complete rhesus macaque *KIR* cluster. This region covered 330 kb on the rhesus macaque reference genome (Mmul_10). Ironically, the *KIR* gene cluster is poorly assembled and annotated on this reference genome, and might not be used as a reference to assemble *KIR* haplotypes.

The rhesus macaque *KIR* cluster was enriched from samples of three animals. One or two flowcells were used to enrich fragments that comprised *KIR* genes from start to end, or connected neighboring *KIR* genes. A range of 2.5% to 36.3% of the total reads had a read-length of 7,000 bp or longer (Table 2). The percentage captured on-target reads ranged from 0.5% to 4.2% (Table 2). The mean coverage of the target region was determined by mapping all reads to the reference genome (Mmul_10), which was complemented with the assembled *KIR* haplotypes as artificial chromosomes, and reached 26X to 91X. The enrichment factor ranged from 128X to 637X. Consensus sequences were generated, which displayed lengths ranging from 7.2 to 20.0 kb. The accuracy of the consensus sequences is estimated by a comparison with exon sequences from the non-human primate KIR database (IPD-NHKIR, release 1.3.0.0), and reached 96.7% to 100% similarity at these coding regions.

**Table 2:** Overview of the total reads, read length, on-target hits, and the coverage of the target region in rhesus macaque samples. *For #3, a single flowcell was used for all different crRNA pools, generating fragments containing a *KIR* gene from start to end, and fragments that span from one *KIR* gene to another.

|  | Gene fragments (flowcell 1) |  |  |
| --- | --- | --- | --- |
|  | #1 | #2 | #3 |
| Total reads | 252981 | 453649 | 2208000 |
| Total reads > 7.000 bp | 25062 | 54772 | 55490 |
| Percentage reads > 7.000 bp | 9.9 | 12.1 | 2.5 |
| Average read length (bp) | 3098 | 3375 | 1978 |
| Average read length (bp) of reads > 7.000 bp | 11258 | 11875 | 9319 |
|  | Gene to gene fragments (flowcell 2) |  |  |
|  | #1 | #2 | - |
| Total reads | 48586 | 20956 | - |
| Total reads > 7.000 bp | 5179 | 7607 | - |
| Percentage reads > 7.000 bp (%) | 10.7 | 36.3 | - |
| Average read length (bp) | 2884 | 7110 | - |
| Average read length (bp) of reads > 7.000 bp | 12226 | 14789 | - |
|  | Total reads (flowcells 1 and 2) |  |  |
|  | #1 | #2 | #3 |
| On-target reads (> 7.000 bp reads) | 1070 | 1005 | 292 |
| Percentage on-target reads | 4.3 | 1.8 | 0.5 |
| Mean coverage target region | 60.8 | 91.8 | 26.8 |
| Enrichment factor | 637 | 404 | 128 |

An allele level resolution allowed the phasing of six rhesus macaque *KIR* haplotypes (Fig. 3). Even with a single flowcell, sufficient coverage was reached to define haplotypes at an allele level resolution (Fig. 3C). The largest haplotype contained 16 *KIR* genes and spanned 280 kb (H15), whereas the shortest *KIR* haplotype encoded five members (H10). A fusion gene, which consists of segments from two distinct *KIR* genes, and that are occasionally generated by chromosomal recombination events, was identified on H14 (*KIR3DL20*030R*) (Fig. 3A). These recombined entities often remain undetected by current genotyping approaches.

**Figure 3.**
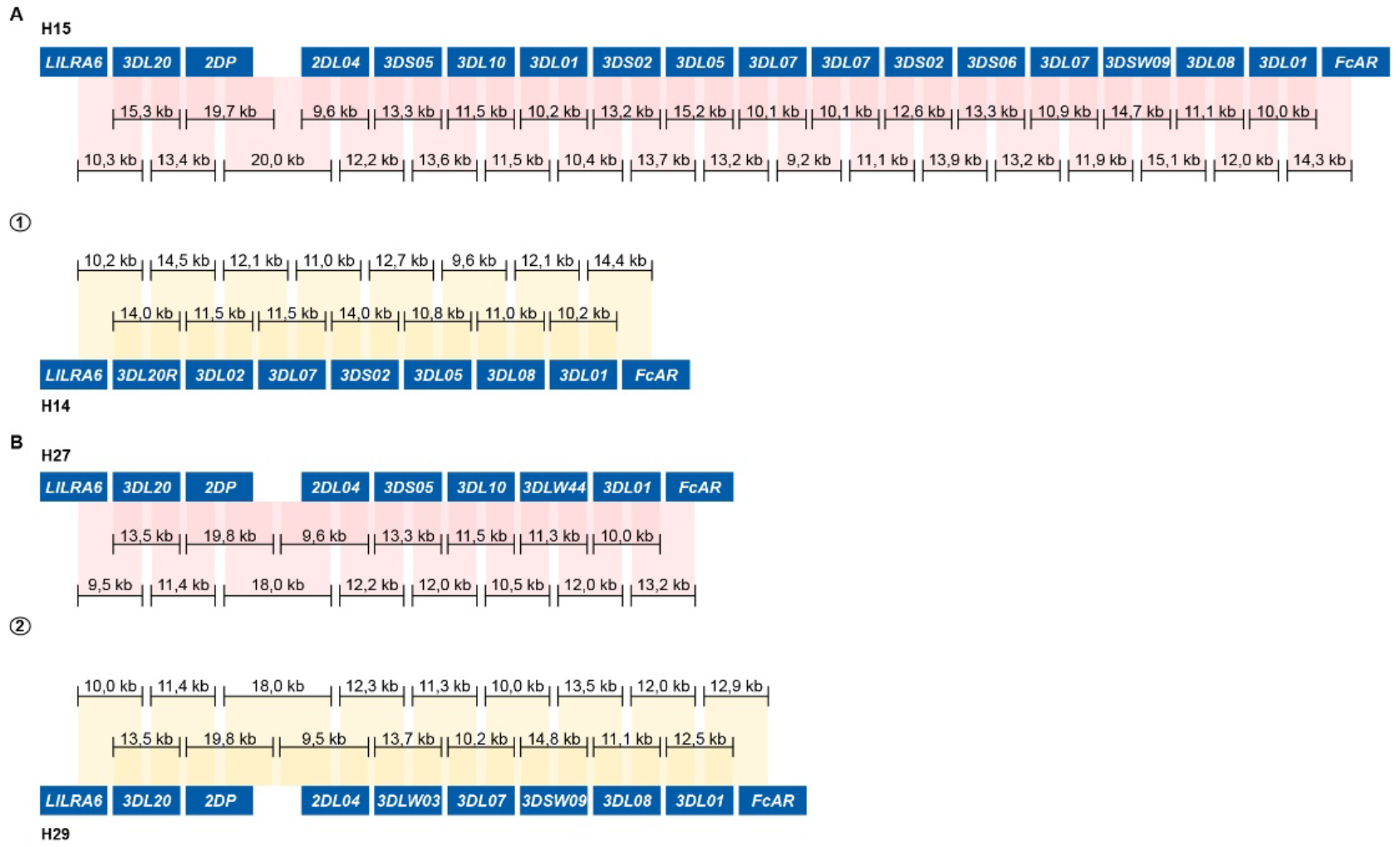
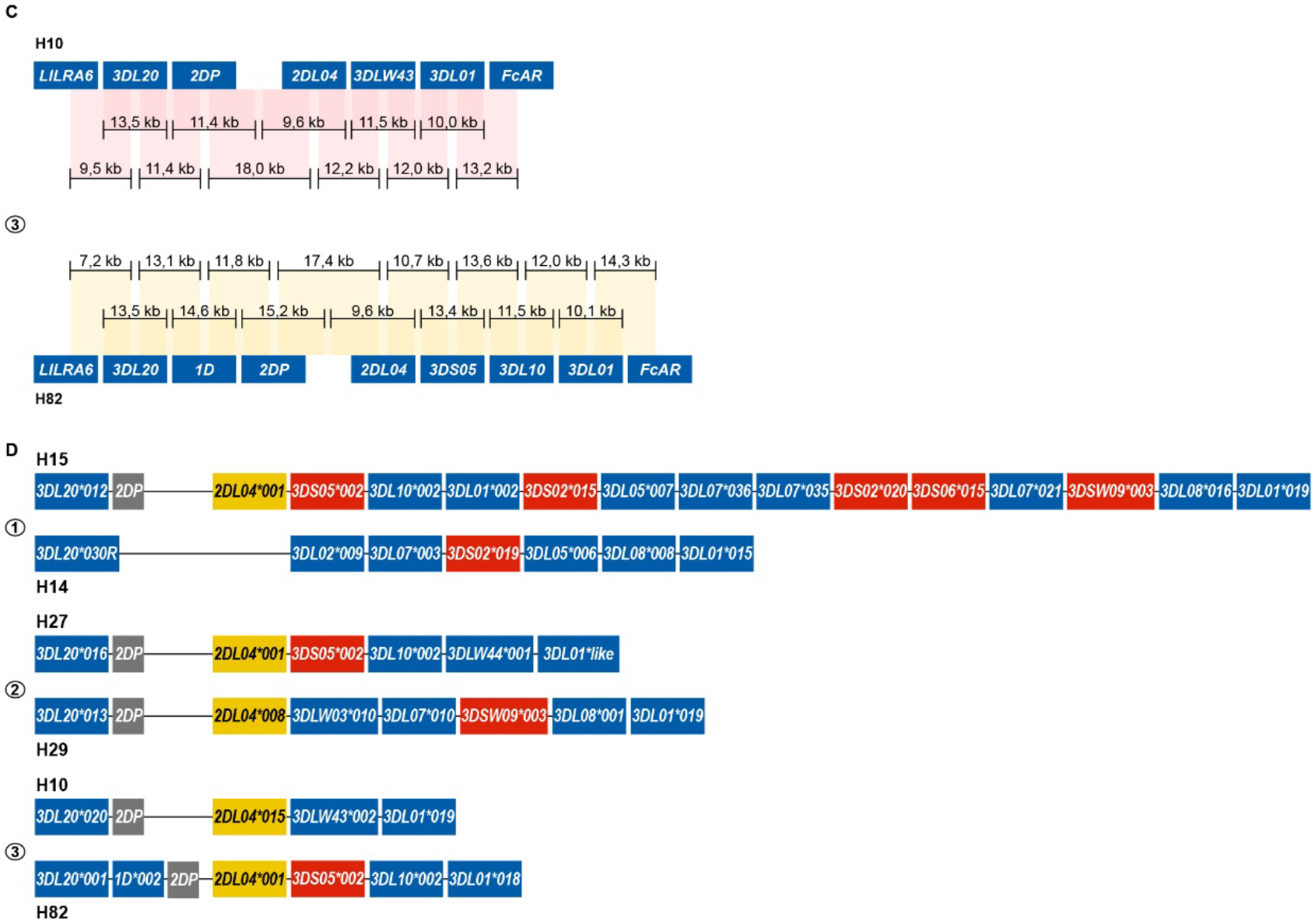
Enrichment and phasing of rhesus macaque *KIR* haplotypes. For rhesus macaque samples (#1, #2, and #3), the *KIR* gene content was determined by a combination of transcriptome and segregation studies [32-34]. Our approach resulted in an accurate annotation of the complex *KIR* regions present in these animals. Consensus sequences (black bars) that covered *KIR* genes from exon 1 to exon 9, or that comprised segments from neighboring *KIR* genes, were assembled. The physical order of *KIR* genes at the haplotype was determined by the extensive overlaps and the alignment with exon sequences from the IPD-NHKIR. The consensus sequence lengths are indicated. Phasing was achieved for all haplotypes, and reflected six different configurations (A, B, C). For each sample, haplotypes were sorted out at an allele level resolution (D). One new allele was defined for *KIR3DL01* (H27), which is indicated with “like”.

### Methylation profile of the multigenic KIR region

Amplification-free enrichment and nanopore sequencing allow the characterization of DNA modification profiles. *KIR* genes display a variegated expression on NK cells and subsets of T cells, which is tightly regulated by methylation of the promotor region (Chan et al. 2003). Therefore, we decided to test whether we could determine methylation signatures for the *KIR* gene region. A high modification likelihood and frequency was determined for the promotor region of all *KIR* genes in the human samples (Fig. 4A). The modification frequency of the complete intergenic region ranged from 62% to 93%, and seems to slightly increase in the proximal promotor adjacent to exon 1. The intergenic regions in the rhesus macaque *KIR* cluster also displayed high likelihood and frequency of modification (Fig. 4B). However, the slight increase of modification frequency at the proximal promotor is not observed. Because all DNA samples were isolated from peripheral blood lymphocytes, the high methylation frequency is in line with scare expression of KIR on most prominent lymphocytes. The true resolution power of this approach would emerge using isolated cell populations or single-cell clones.

**Figure 4.**
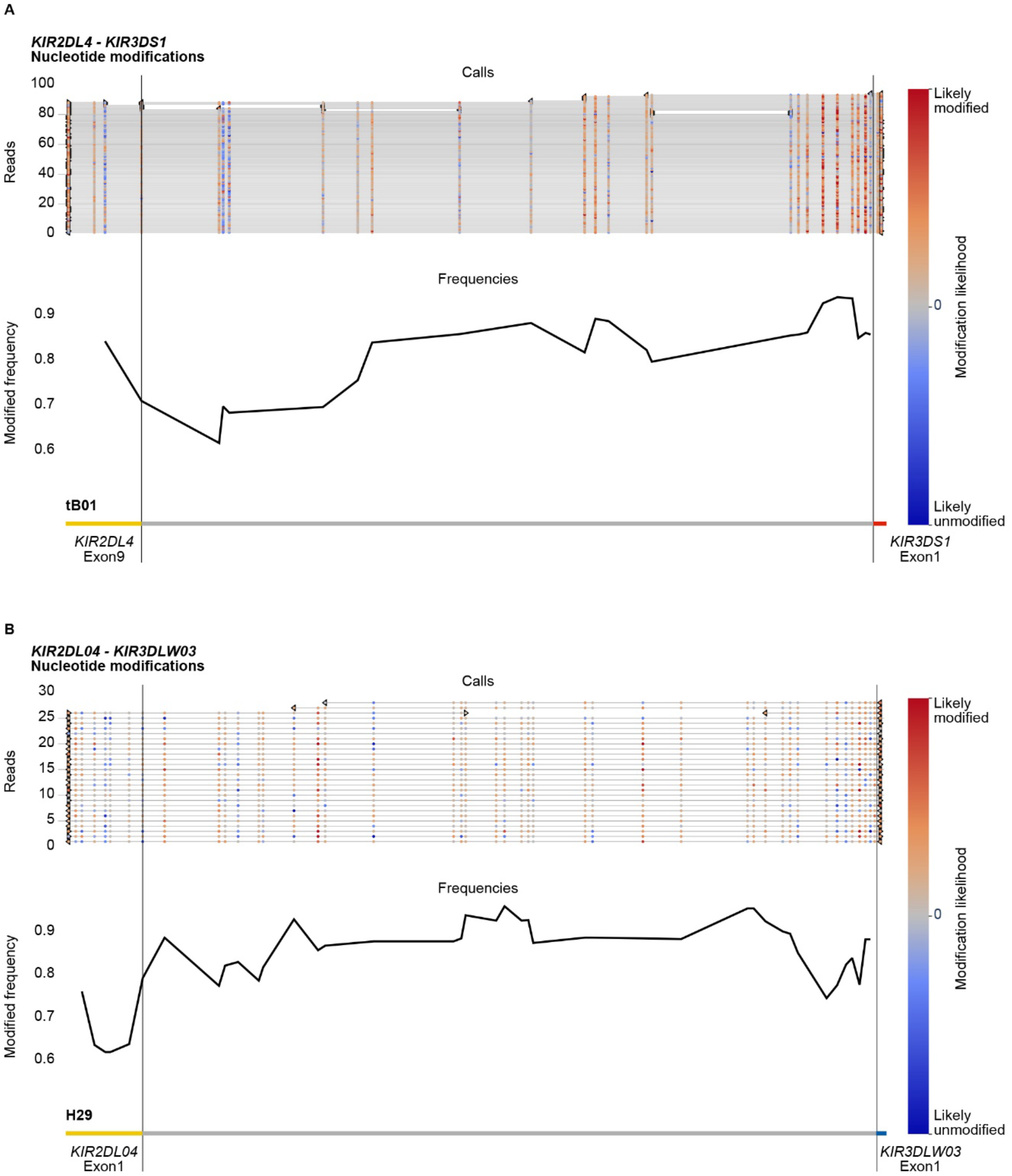
DNA modification profiles. DNA modification profiles are displayed for one human (sample #2) (A) and one rhesus macaque (sample #2) (B) KIR gene. The epigenetic predictions were calculated with up to 100 randomly selected reads per enriched KIR segment. Indicated are the modification likelihoods (red, high; blue, low), the modification frequency, which ranges from 0 to 1, and the annotation that is based on the (complemented) reference genomes (HG38 and Mmul_10). Plots were generated by Methplotlib (De Coster et al. 2020). The *KIR* promotor region, which is approximately 300 bp in front of exon 1 (Chan et al. 2003), is highly modified for human KIR3DS1 (A) and macaque KIR3DLW03 (B), with modification frequencies ranging from 85 to 95%. These two regions are representative for all other *KIR* gene regions studied in our human and rhesus

## Discussion

Most multigenic regions are subjected to concerted evolution, in which species-specific gene duplications, deletions, and recombination events shape a genetic cluster. Immune gene families, such as KIR and MHC, diversified under selective pressure by a continuous arms race with pathogens, and individuals might benefit from extensive variation at the population level (Nei and Rooney 2005). The high gene content diversity and sequence similarity challenges rapid genomic characterization of these multigenic families.

We make available a rapid enrichment and characterization approach for complex multigenic regions, which overcomes many of the limitations of short-read or whole-genome sequencing. Extensive diversity in gene content of multigene families was resolved by targeting long overlapping genomic fragments that covered genes from start to end, or that contained segments of neighboring genes. Efficient targeting was based on using generic crRNAs, which yielded a high median coverage at the region of interest, and allowed the generation of relatively accurate consensus sequences. Most inaccuracies involved homopolymers, here defined as a stretch of three or more identical bases, which simultaneously impact the current measured by the nanopore. The consensus sequence accuracy was, however, sufficient to define alleles and to phase complex *KIR* haplotypes. Only at large identical stretches in which overlapping fragments lack allelic variation might this technique hamper the phasing of haplotypes (Fig. S2). However, the continuous improvement of library preparation chemistry, base-calling algorithms, and post-sequencing correction software might eventually increase the length of enriched fragments and the accuracy of nanopore sequencing.

The target coverage was sufficient for the characterization of *KIR* haplotypes at the allele level resolution, but the enrichment performance displayed some deviations in the different samples (Tables 1 & 2). The variance in total read numbers and read length is likely to be affected by flowcell variations, gDNA sample quality, and short read clean-up efficiency during library preparation. Despite the deviations in read counts, the on-target coverage was sufficient, even when only a single flowcell was used (Table 2, #3; Fig. 3C). This ensured highly accurate and overlapping consensus sequences that resolved complete human and macaque *KIR* haplotypes at an allele level resolution (Fig. 2D & 3D).

The high modification frequency of the *KIR* promotor regions (Fig. 4) indicates low levels of expression in the characterized samples (Chan et al. 2003). This is likely explained by a selective and variegated expression of KIR receptors, which are only defined on the cell surface of NK cells and subsets of T cells. Our HMW DNA samples were isolated from whole blood, which includes only a small fraction of KIR-positive cells. Nevertheless, the epigenetic modification of members from highly complex multigenic families could be determined using our approach. This might provide biological and diagnostic insights into the role of these complex systems in health and disease.

Members of multigenic families often encode components of essential immune responses, and exhibit extensive copy number variation and allelic polymorphism. The continuous diversification and selection of these genes enables adaption to pathogens, but might also generate candidates that enhance susceptibility to immune-related disorders. For instance, many HLA and KIR allelic polymorphisms are known to have an impact on human health, susceptibility or resistance to disease, or may affect graft survival success in transplantation biology (Hiby et al. 2004; Farag et al. 2006; Bashirova et al. 2011). Many of the relevant SNPs are mapping in coding regions, whereas others are not, and may impact, for instance, gene (expression) regulation and alternative splicing potential. This enrichment technique fosters a cost-efficient and rapid strategy to characterize multigenic families at an allele level. Insights regarding complex immune clusters might provide a comprehensive perspective on biological interpretations, and lift SNP disease association studies from the allele to the haplotype/region level, in which all polymorphisms are considered.

## Materials and methods

### Cells and genomic DNA extraction

Human buffy coat samples from healthy donors were obtained from the Dutch blood bank (Sanquin, the Netherlands). Rhesus macaques with a characterized KIR transcriptome were selected from the self-sustaining colony housed at the Biomedical Primate Research Centre (BPRC) (Bruijnesteijn et al. 2018; Bruijnesteijn et al. 2020b). Heparin whole blood samples from these animals were obtained during annual health checks. PBMCs were isolated from human buffy coats and rhesus macaque heparin samples.

High-molecular-weight (HMW) gDNA was isolated from human and rhesus macaque PBMC samples (± 7 × 10^6^ cells), using the Circulomics Nanobind CBB Big DNA Kit (Circulomics, NB-900-001-01) and following the manufacturer’s instructions. The concentration and purity of the gDNA samples were determined using a Nanodrop and a Qubit platform. The (HMW) gDNA fragment length was determined by pulsed field gel electrophoresis (PFGE) in reference to a lambda PFGE ladder.

### Designing guiding crRNAs and constructing RNPs

For both humans and macaques, sets of generic and specific CRISPR RNAs (crRNA) were designed within the *KIR* gene cluster and the flanking *LILR* and *FCaR* genes by using Benchling, a freely available online software tool (Benchling 2020). The crRNAs have a guiding length of 20 bp, and are designed in front of a protospacer adjacent motif (PAM) sequence “NGG”, in which “N” represents any nucleotide base. The software tool provides on- and off-target scores for specific crRNAs. The on-target scores are based on optimized calculations from Doench and colleagues, with the higher score indicating the better crRNA target binding (Doench et al. 2016). The off-target scores reflect the specificity of the crRNAs, and were based on different builds of the human (NCBI36, GRCh37, GRCh38) and macaque (MMUL_1, Mmul_8.0.1) reference genomes (Hsu et al. 2013). CRISPR RNAs with high off-target scores were considered specific for one *KIR* gene. Relatively low off-target scores (ranging from 3 to 60) were recorded for crRNAs that potentially target multiple *KIR* genes and were included in the panel as putative generic crRNAs. In total, 54 and 45 custom crRNAs were selected to enrich the human and macaque *KIR* gene cluster, respectively (Sup. Table 1 and 2) (IDT, custom Alt-R® CRISPR-Cas9 crRNA). Sets of different crRNAs were pooled based on the different strand-directed orientations (sense versus anti-sense) to avoid cleavage and the sequencing of unintended short on-target fragments. The pools were defined to generate DNA fragments that comprise *KIR* genes from exon 1 to exon 9, or fragments that connect neighboring *KIR* genes (Sup. Table 1 and 2). The pooled crRNAs were mixed with trans-activating crRNAs (tracrRNA) (IDT, #1072534) in a 1:1 ratio and further diluted in Duplex Buffer to a final concentration of 10 μM. The crRNAs and tracrRNA were annealed by heating the duplex solution for 5 min at 95 °C, followed by cooling to room temperature (RT) on a benchtop. To subsequently construct the Cas9 ribonucleoprotein particles (RNPs), the crRNA-tracrRNA duplexes were assembled with HiFi Cas9 endonuclease (IDT, #1081060) in 1x NEB CutSmart Buffer (NEB, #B7204S) at a total volume of 30 μl by incubating the solution for 30 min at RT. The Cas9 RNPs were stored until use at 4 °C for up to a week.

### Cas9-mediated target enrichment and Oxford Nanopore sequencing

Throughout the protocol, unintended fragmentation of gDNA will decrease the capturing efficiency and enrich off-target fragments. Therefore, samples should be handled with care and processed with wide-bore pipette tips. Input gDNA (5-10 μg) was resuspended in 10x NEB CutSmart Buffer (8:1) and dephosphorylated by incubation with Quick calf intestinal phosphatase (CIP) (NEB, # M0525S) at 37 °C for 20 min, followed by heating at 80 °C for 2 min to deactivate the enzyme (Fig. S1). After the sample returned to room temperature, the dephosphorylated gDNA (30 μl) was gently mixed with a Cas9 RNP pool (10 μl), 10 mM dATP (1 μl), and Taq polymerase (1 μl), followed by incubation at 37 °C for 60 min, then 72 °C for 5 min, and hold at 4 °C. Ligation buffer (20 μl) and sequencing adaptors (5 μl) from the Ligation Sequencing Kit (ONT, #LSK109) and Quick T4 DNA Ligase (10 μl) (NEB, M2200S) were added to the cleaved and dA-tailed gDNA sample, followed by an incubation of 60 min at RT. Adapter-ligated samples from the same individual, which were treated with different crRNA sets, were pooled, and diluted (1:1) in TE buffer. The excess of adaptors and short DNA fragments were removed using 0.3x AMPure XP Beads (Beckman Coulter, #A63881), which were washed twice on a magnetic rack with Long Fragment Buffer (ONT, #LSK109). The beads were eluted in 15 μl Elution Buffer (ONT, #LSK109). A sequencing library was prepared by adding 37.5 μl Sequencing Buffer and 25.5 μl Loading Beads (ONT, #LSK109) to the processed DNA sample. Eluted samples were sequenced on a R9.4.1 flowcell using an Oxford Nanopore MinION device. Prior to sequencing, the flowcells were primed according to the manufacturer’s instructions using the Flow Cell Priming Expansion Pack (ONT, #LSK109). After 24 hours of sequencing, flowcells with over 500 active pores remaining were washed and reloaded with a second enriched library of the same gDNA sample according to the manufacturer’s instructions using the Flow Cell Wash Kit (ONT, #EXP-WSH003).

### Sequence data analysis

Basecalling and read quality assessment (min qscore 7) were performed using Guppy V3.4.1 software on a Linux platform that utilized a GeForce RTX 2080 Ti graphics processing unit (GPU). Basecalled reads were imported into Geneious Prime software (v.2020.1.2) for further analysis. Exon libraries, including sequences of exons 3, 4, and 7 from the human and rhesus macaque KIR databases (IPD-KIR, release 2.9.0; IPD-NHKIR, release 1.3.0.0), were used as references to map the Nanopore reads into contigs based on similarity. Each on-target contig resembled a specific *KIR* gene or a pair of neighboring *KIR* genes

For each human individual, the haplotype configurations (e.g., cA01-tA01) could be largely deduced from the read contigs generated by exon library mapping. On the basis of this knowledge, the reads of a single flowcell were re-mapped to the complete human reference genome (HG38), complemented with the particular *KIR* haplotype configuration references, using minimap2 (version 2.17). Consensus sequences that covered *KIR* genes from start to end, or that comprised segments of neighboring *KIR* genes, were generated from the alignments based on 65% nucleotide similarity. The consensus accuracy was determined by comparison to the reference haplotype. The on-target coverage was resolved for each individual by mapping all reads with a read length above 7,000 bp to the human reference genome (HG38) using minimap2 in Geneious software. The enrichment factor was determined as the ratio of on- and off-target mean coverage, and reflects the enrichment efficiency. For each rhesus macaque sample, reads from the on-target exon contigs were re-mapped to the contig consensus sequence using minimap2. Additional re-mapping steps were performed using the generated consensus sequences to optimize accuracy and diminish homopolymer errors. To further optimize these consensus sequences, shorter on-target reads (< 7,000 bp) were included for consensus calculations. The eventual consensus sequences were generated based on 65% nucleotide similarity. The consensus accuracy was estimated by the alignment with exon references available from previous KIR transcriptome studies (IPD-NHKIR, release 1.3.0.0). On-target coverage was defined by mapping all reads with a length of 7,000 bp or more to the rhesus macaque reference genome (Mmul_10), which was complemented with the appropriate assembled *KIR* haplotypes, using minimap2 in Geneious software.

### DNA modification profiles

DNA modifications, such as methylation, were called using Guppy V4.0.12 software with the dna_r9.4.1_450bps_modbases_dam-dcm-cpg_hac.cfg configuration and fast5_out flag. Multi-read fast5 files were converted to single-read files using multi_to_single_fast5 provided by ONT. The basecalled reads were mapped to a reference genome using minimap2, and subsequently called for modifications using Nanopolish. The methylation likelihood and frequencies were visualized by Methplotlib (De Coster et al. 2020), using an annotated reference genome. When the reference genome did not contain the appropriate *KIR* genes, like Mmul_10 for all macaque individuals, methylation calls were annotated using a modified reference genome, which was complemented with the newly assembled haplotypes as artificial chromosomes.

## Acknowledgements

We thank D. Devine for editing the manuscript and F. van Hassel for preparing the figures.

## Data availability

All raw Nanopore files and processed sequencing data generated in this study have been submitted to the European Nucleotide Archive (ENA) (https://www.ebi.ac.uk/ena/browser/home) under accession number PRJEB43311.

## Supplemental figure legends

**Figure S1. Library preparation of the targeted enrichment protocol.** A schematic overview of the library preparation to enrich a target region (blue). Freshly isolated HMW genomic DNA was dephosphorylated by incubation with CIP for 20 min at 37 °C. Subsequently, selected pools of RNPs (Sup. Table 1 and 2) were added to aquilots of the dephosphorylated gDNA, and incubated for 60 min at 37 °C in the presence of Taq polymerase and dATP, followed by incubation for 5 min at 72 °C. The available phosphate groups at the terminus of the targeted region were ligated to Nanopore adaptors during incubation with T4 ligase for 60 min at room temperature (RT). The adapter-ligated aliquots were pooled, followed by clean-up with 0.3X AMPure beads. The eluted sample was loaded on a flowcell for Nanopore sequencing.

**Figure S2. A SNP desert at the *KIR2DL1* gene.** Human individual #3 shared an identical *KIR2DL1* allele at both haplotypes, which located a 25 kb SNP desert. The lack of reads that span the complete SNP desert hampered the phasing of *KIR2DL1*.

